# PfaSTer: A ML-powered serotype caller for *Streptococcus pneumoniae* genomes

**DOI:** 10.1101/2022.11.30.518579

**Authors:** Jonathan T. Lee, Xingpeng Li, Craig Hyde, Paul A. Liberator, Li Hao

## Abstract

*Streptococcus pneumoniae* (pneumococcus) is a leading cause of morbidity and mortality worldwide. Although multi-valent pneumococcal vaccines have curbed the incidence of disease, their introduction has resulted in shifted serotype distributions that must be monitored. Whole genome sequence (WGS) data provides a powerful surveillance tool for tracking isolate serotypes, which can be determined from nucleotide sequence of the capsular polysaccharide biosynthetic operon (*cps*). Although software exists to predict serotypes from WGS data, their use is constrained by the requirement of high-coverage Next Generation Sequencing (NGS) reads. This can present a challenge in so far as accessibility and data sharing. Here we present PfaSTer, a method to identify 65 prevalent serotypes from individual *S. pneumoniae* genome sequences rather than primary NGS data. PfaSTer combines dimensionality reduction from k-mer analysis with machine learning, allowing for rapid serotype prediction without the need for coverage-based assessments. We then demonstrate the robustness of this method, returning >97% concordance when compared to biochemical results and other *in-silico* serotypers. PfaSTer is open source and available at: https://github.com/pfizer-opensource/pfaster.

## Introduction

*Streptococcus pneumoniae* (pneumococcus) presents a major concern to public health, being a common cause of lower respiratory tract infections and pneumonia [1, 2]. Pneumococcal disease is a particular threat to the elderly, largely due to a high mortality risk when contracting pneumonia [1, 3]. Pneumococcal conjugate vaccines (PCVs) can be used to prevent disease [4, 5] by affording protection against common circulating serotypes. In *S. pneumoniae*, serotype is defined by the structure of a capsular polysaccharide and the genes that direct biosynthesis of the polysaccharide encoded at the capsular polysaccharide synthesis (*cps*) operon [6]. To date, over 95 pneumococcal serotypes carrying unique *cps* sequences have been identified [7], with a fraction of these found to be prevalent in global populations [7]. As the capsular polysaccharide serves as the target of PCVs [4], surveillance of emerging strains through serotyping is important for monitoring efficacy against circulating strains and the development of new multi-valent vaccines [8].

Traditionally, pneumococcal serotyping is performed using serotype-specific monoclonal antibody reagents, either through the Quellung reaction or latex agglutination [9]. While held in high regard, such methods are expensive and laborious [9, 10]. Antibody tests are also often unable to differentiate closely related serotypes [9, 11], and visual assessment of agglutination results are susceptible to subjective interpretation. Furthermore, the need for cell cultures presents a physical barrier for replicating results between research groups. As an alternative, automated pipelines for predicting serotypes from Next Generation Sequencing (NGS) data have been developed. Since 2016, PneumoCaT, SeroBA, and more recently SeroCall, have been utilized to effectively identify serotypes *in-silico* [10, 12, 13]. While their underlying algorithms differ, these methods all utilize the same input: raw NGS data from the *cps* locus and a reference *cps* database for different serotypes. By leveraging an abundance of NGS reads, these applications provide robust predictions of the *cps* sequence and therefore the *in-silico* serotype.

While a powerful resource, high-coverage NGS data can be unwieldy and computationally intensive to work with. Furthermore, such data is not always readily available to researchers. For instance, the PubMLST [14] microbial database contains, to date, over 30,000 pneumococcal genomes from submissions around the globe. Many of these assembled genomes lack accompanying NGS data sources and would be incompatible with the previously described serotyping tools.

We developed the pneumococcal FASTA serotyper (PfaSTer) to address the need for *in-silico* serotyping when constrained to working with assembled or aligned genome sequences. PfaSTer identifies k-mers at the *cps* locus associated with each serotype, which are utilized in machine learning for prediction (Fig 1). Using a validated dataset of >2,000 pneumococcal isolates, we show that PfaSTer is both a fast and highly accurate serotype caller, with predictions comparable to both serological results and other computational methods.

**Fig 1:**
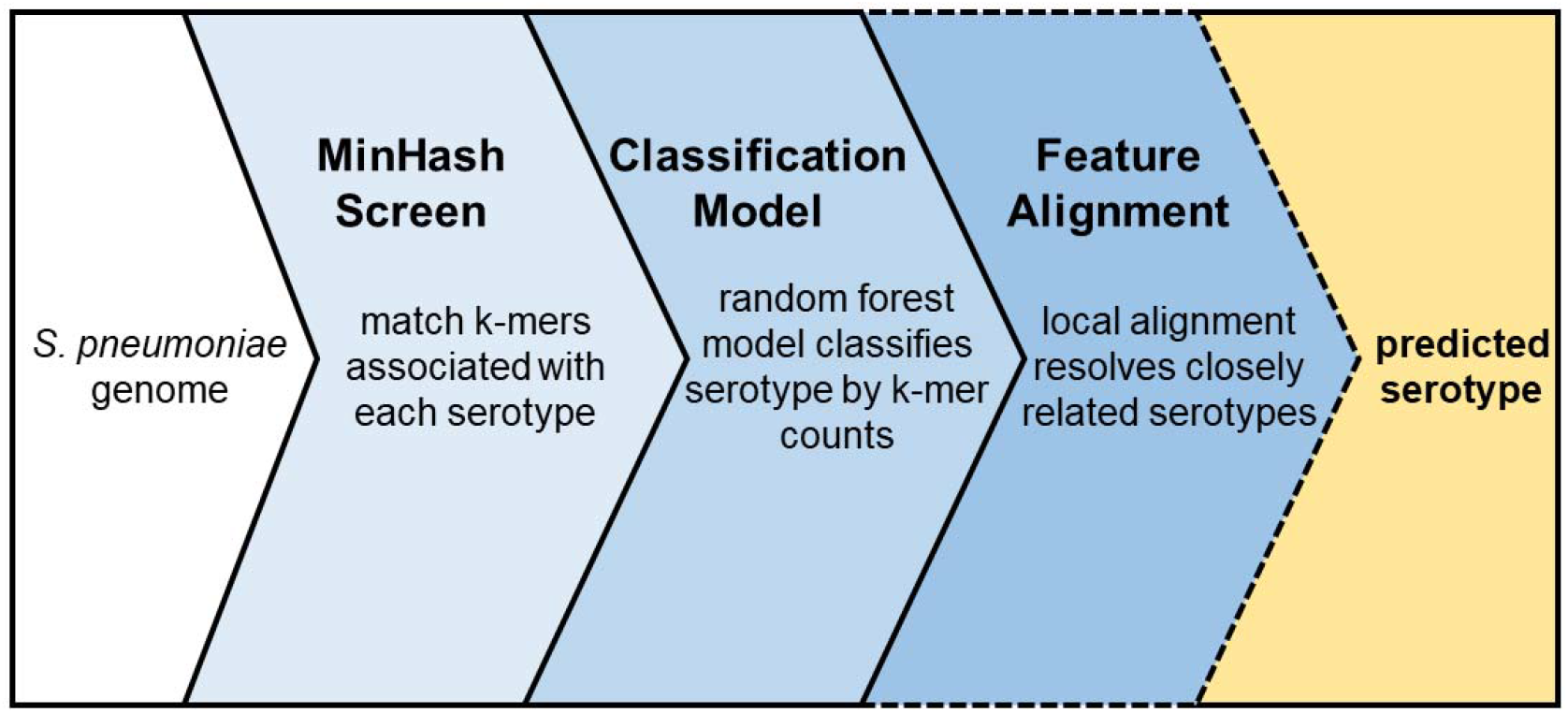
PfaSTer workflow. PfaSTer takes an aligned or assembled *S. pneumoniae* genome sequence (FASTA format) as an input. A MinHash screen for k-mers associated with each reference serotype is first performed. The number of k-mers matched to each reference is then passed to a Random Forest classifier to assign a predicted serotype. In cases where the model is unable to discern closely-related types, alignment is performed to identify serotype-defining features.

## Materials and Methods

### Data sources

Training data for PfaSTer was obtained from the Sanger Institute Pathogenwatch platform (pathogen.watch) in the form of de-novo assembled genomes for isolates spanning 65 different serotypes (Table S1). For validation, sequences were obtained from the NCBI sequence read archive. Accessions for these data can be found in (Table S2).

### Mash sketch creation

Reference *cps* sequences (previously published and utilized by PneumoCAT [12] and seroBA [10]) were used to develop a MinHash sketch [15] of 65 serotypes. A sliding window (k-mer) of 70 nucleotides was used to scan each *cps* sequence, with each k-mer converted to a 128 bit integer using MurmurHash3 (v3.0.0). To account for bidirectionality, both the forward and reverse complemented k-mer were considered and the lexicographically smaller sequence used for hashing. The k-mers corresponding to the 1,000 smallest integer values for each serotype were saved to the sketch.

### Model training and probability thresholding

A Mash screen [16] was performed for 4,019 pneumococcal genomes using the previously described sketch. Each hash of 70 base pair k-mers in a sliding window across the genome sequence was compared to those in the reference sketch, and matching k-mers recorded. The total number of k-mers matched for each serotype were then saved and used as features to train a Random Forest classifier using the R tidymodels package (v0.1.2). To account for class imbalance due to differences in serotype prevalence, overrepresented serotypes were down sampled to no more than 200 cases for training. Initial model performance was measured using a grid search and the average accuracy across 2000 internal cross-validations. The model was then ported to python using the sklearn package (v1.1.1). Hyperparameter tuning was performed using a grid-search, with optimal parameters found to be 300 estimators, 10 features per estimator, and 4 samples to split branches. Model performance was re-calculated and reported using the average accuracy across 200 internal cross-validations.

To limit errant predictions, the model-computed probability of both correct and incorrect predictions was recorded for each serotype based on the training dataset in cross-validation. For each sample, the serotype with the highest prediction probability was saved and noted as correct or incorrect classification compared to their labeled serotype. The probability distributions of correct and incorrect classifications were used to fit a generalized linear model with a binomial distribution for each of 17 serotypes. For cases where the two distributions did not overlap, a minimum probability threshold was determined as (ln(p/(1-p)) - b_0_)/b_1_, where b_0_ is the fitted intercept, b_1_ the slope, and p = 0.05. For cases where the distributions did overlap, the minimum threshold was calculated using the upper limit of the one-side 95% confidence interval of the incorrect classification distribution.

### Feature alignment for closely related serotypes

Reference sequences for *wciZ* (serotype 15B), *wciX* (serotype 18C), and *wciG* (serotype 35B) were obtained from annotated genomes at NCBI (accessions CR931664, CR931673, and KX021817, respectively). BLASTN [17] was used to obtain the sequence of the corresponding gene for each serotype, and the resulting reading frame was assessed for presence of a premature stop codon.

### Validation with an external dataset

A collection of short-read sequencing data for 2,065 UK isolates originally from Public Health England was used for validation. Reads were de-novo assembled to genome sequences using SPAdes (v3.14.0, - isolate mode) [18] and serotypes predicted using PfaSTer. Isolates that were previously labeled through latex agglutination [10] to be non-typeable, or serotypes not supported by PfaSTer, were excluded from calculations. This resulted in validation against 2,026 samples (Table S2). PfaSTer predicted serotypes were compared to latex agglutination results as well as calls made by both PneumoCaT and SeroBA – previously reported in [10] (Note S1).

## Results

We sought to develop a method for predicting pneumococcal serotypes relying only on minimal data in the form of consensus genome sequences. To this end, we first applied the MinHash (Mash) algorithm, a dimensionality-reduction technique that can effectively compress up to entire genome sequences to a small collection (or *sketch)* of several thousand sub-sequences (k-mers) [15]. As the capsular polysaccharide is encoded at the *cps* operon, we started by performing a Mash Screen [16] comparing >4,000 pneumococcal genomes against a k-mer sketch of each serotype’s *cps* locus. The number of matched k-mers to each serotype was then used as features to train a Random Forest classifier. This method predicts the pneumococcal serotype based on the collective voting of hundreds of decision tree estimators, each trained on a bootstrapped set of the >4,000 training samples.

Through internal cross-validation, we found the resulting model yielded a median accuracy of 97.8% in our training data. To account for misclassification from low-confidence predictions, we recorded the prediction probabilities returned by the Random Forest model during cross-validation and calculated the probability distributions of correct and incorrect serotype calls (Fig S1). We then set thresholds based on the 95% confidence intervals, flagging serotype predictions below these values as low-confidence. Following this addition, most remaining misclassifications resulted from closely related serotypes, which could not be distinguished using the Mash screen results due to a high density of shared k-mers (Fig S2). In particular, the serotype pairs 15B/C, 18B/C, 24B/F, and 35B/D had a higher rate of incorrect serotype calls compared to other types during cross-validation (Table S3). While the genetic cause of the 24B and 24F capsular polysaccharides has previously been hypothesized and studied [6, 19], the exact mechanism underlying their differing polysaccharide structures is still unclear. As we cannot reliably distinguish serotype 24B from 24F at this time, PfaSTer reports Serogroup 24 when either of these types is predicted by the model. In contrast, modifications that inactivate genes that code for O-acetyltransferases (*wciZ* for 15B/C, *wciX* for 18B/C, and *wxiG* for 35B/D) [20–22] impact polysaccharide structure and serotype designations. These modifications can include in/dels as well as SNVs leading to frame shifts and/or premature stop codons. Unfortunately, subtle and heterogeneous modifications that inactivate a step in polysaccharide biosynthesis and therefore polysaccharide structure are generally not detectable with the Mash screen technique.

To overcome this challenge in classifying 15B/C, 18B/C, and 35B/D isolates, we added a local alignment stage when one of these serotypes is predicted by our model. This step searches the corresponding acetyltransferase for premature termination that would inactivate the protein. By applying this check, we were able to successfully assign each isolate to the correct serotype.

As final validation, we applied the PfaSTer prediction pipeline to 2,026 isolates previously evaluated using the *in-silico* serotyping tools PneumoCaT [12] and SeroBA [10], both of which utilize NGS read data as inputs. Compared to results from latex agglutination, PfaSTer showed 97.09% concordance in its serotype predictions (Fig 2, Table S2). This is similar to the ~98% concordance previously reported by Epping et al using PneumoCat and SeroBA [10]. Furthermore, serotype calling by PfaSTer was in high concordance with the other computational methods, returning the same serotype as PneumoCaT in 97.97% of cases and SeroBA in 98.47% of cases (Fig 2, Table S2). PfaSTer also demonstrated an extremely rapid runtime during this benchmarking, with all 2,026 samples completed in under 2 hours on a 36 cpu Amazon EC2 c4 instance running 8 parallel processes.

**Fig 2:**
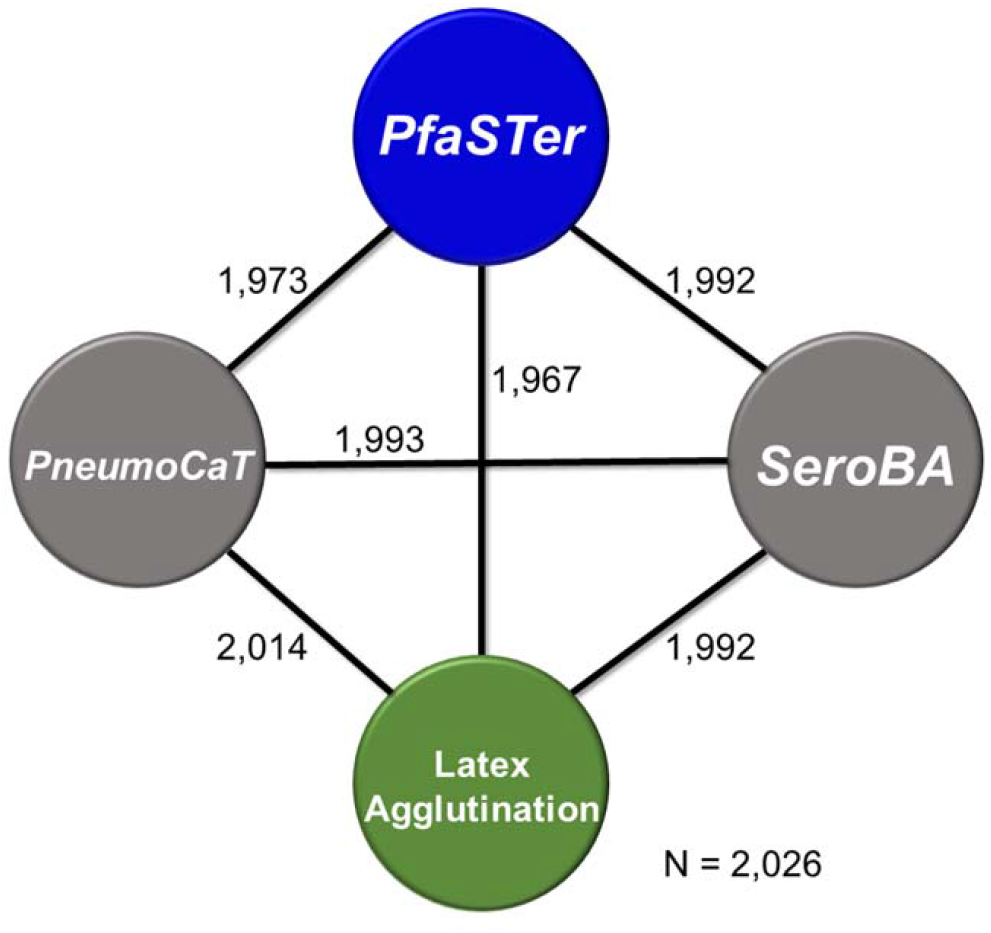
Concordance between serotyping methods for a validated dataset. Number of isolates (out of 2,026) returning the same serotype when tested using PfaSTer, latex agglutination, and two other *in-silico* serotyping tools

Among the isolates used for comparison, 17 were not typed by PfaSTer due to prediction probabilities falling below our computed thresholds. For cases where PfaSTer predicted a serotype with high confidence, most disagreement with other typing methods occurred with 15C and 35D designations (Table S2). There were 19 instances where results from latex agglutination, PneumoCaT, and SeroBA differed from not only PfaSTer, but also one-another when calling the serotype as 15B or 15C. PfaSTer also identified six isolates as serotype 35D, which were identified as 35B by PneumoCaT and latex agglutination. As an additional validation, we saved the *wciG* alignments for the six 35D predictions, and the *wciZ* alignments for five randomly selected 15C predictions that differed from latex agglutination. We then reviewed the resulting protein sequences for truncation. In all cases, a premature stop was indeed observed in the corresponding O-acetyltransferase (Fig S3). These isolates are therefore expected to express 15C or 35D capsular polysaccharide, as predicted by PfaSTer.

## Discussion

We have developed an efficient tool for rapid *in-silico* serotyping of *S. pneumoniae* from assembled genome sequences. This method uses a single-pass k-mer screen and a machine learning model to predict the *S. pneumoniae* serotype without needing to access raw NGS data. While a targeted alignment step is included to resolve a small subset of serotype-specific features (a limitation shared among serotyping pipelines [10, 12]), high density read data is not needed, in contrast to other published tools.

A major challenge in developing PfaSTer was establishing confidence in the serotype predictions when constrained to a single genome sequence. While other *in-silico* algorithms designed to assign serotype utilize per-base or per-k-mer coverage to generate confidence, this information is unavailable when working with a single consensus genome. Although the Mash screen results estimate sequence similarity to each serotype, they do not provide any statistical power on their own. This challenge was addressed at the machine learning step by leveraging the innate properties of the Random Forest model. As the model consists of an ensemble of decision trees, prediction probability can be estimated as the proportion of trees agreeing on the serotype [23]. Using these values, thresholds were established to flag low-confidence serotype predictions. These probability estimates are also provided to users of PfaSTer to support their own decision making.

Of the >2,000 isolates used to validate PfaSTer performance, a small fraction exhibited discordance with other serotyping methods. This included 17 samples that did not return a serotype due to low-confidence prediction. Of note, 5 of these isolates were unable to be typed with other *in-silico* pipelines or were reported as a mixture of serotypes (Table S2). This suggests that predictions computed at low probability may be caused in-part by low quality sequencing data.

Discordance was also noted in instances when PfaSTer identified mutations that predict the derived serotypes 15C and 35D. In those samples latex agglutination called the serotype as 15B or 35B, respectively. Notably, each of samples with a lack of 15B/15C concordance mapped to mutations in the same region of *wciZ* at a TA-tandem repeat that has been shown to slip and cause indels during replication (Fig S4) [24, 25]. As repeated frameshifts can convert the serotype between 15B (complete *wciZ* gene and intact O-acetyltransferase open reading frame) and 15C, the serotype can switch over time. [12, 24]. Additionally, as antibodies against 15B have been shown to cross-react with 15C polysaccharide [20, 25], mislabeling could occur when typing with antisera. In contrast to 15C, 35D-causing mutations were more widely distributed across *wciG*, causing premature termination at different positions along the protein coding sequence (Fig S3B). PneumoCaT does not appear to support the identification of serotype 35D, only able to provide a 35B assignment for the samples included in this study. Like the output from PfaSTer, the SeroBA tool also recognized this subset of isolates as serotype 35D.

Although both fast and powerful, serotype assignment using PfaSTer has certain restrictions. As PfaSTer relies on a supervised learning model for prediction, enough cases must be available for training. While over 95 pneumococcal serotypes have been recorded [26], certain serotypes are more prevalent than others throughout the world. As a result, PfaSTer prediction is limited to 65 types due to a shortage of available genomes for rare serotypes. From recent studies, commonly collected serotypes shared across the US, Europe, and Asia include 1, 3, 6A, 6B, 14, 18C, 19F, and 23F, with other serotypes identified at lower frequency [27–31]. Unsurprisingly, these prevalent serotypes are all included in the pneumococcal conjugate vaccine (PCV) formulations of PCV13 [32] and PCV20 [8]. To support continual estimation of vaccine coverage, the commonly circulating serotypes in these PCV formulations are all supported by PfaSTer. As the serotype landscape changes over time, and genomes of new isolates are made available, the number of serotypes predicted by PfaSTer may rise.

By relying on an assembled genome, PfaSTer also has reduced functionality for mixed samples compared to some alignment-based serotype tools. For instance, the SeroCall [13] tool can identify both major and minor serotypes in mixed sequencing data by aligning sequencing reads to multiple references. While PfaSTer does not support prediction for assembled metagenomes, the presence of each serotype can potentially be inferred from the density of k-mers present in the Mash screen step [16]. Future developments on PfaSTer could address this feature more directly.

As global monitoring and sequencing of *S. pneumoniae* continues, PfaSTer provides a means to leverage portable, but previously underutilized, genome sequences for data sharing and serotype tracking. Such surveillance efforts could have important impact on understanding the spread of *S. pneumoniae* and influence future vaccine design for combatting pneumococcal disease. Finally, this method may have applications suitable for typing of other microbial species beyond *S. pneumoniae*.

## Data Summary

PfaSTer is open source and available for Linux on Github under Apache License v2.0 at https://github.com/pfizer-opensource/pfaster Accession numbers for sequencing data are listed in the supplementary material.

## Acknowledgements

The authors would like to thank Charles Tan for his assistance in statistical modeling, and Scott Perrin for digital help in deploying PfaSTer. We also thank Varun Raghuraman for his efforts testing the installation and running of PfaSTer, Zhenghui Li and Andy Weiss for critical reading of the manuscript, and Christina D’Arco for scientific writing assistance.

## Author Contributions

All authors met ICMJE criteria for authorship and participated in the study design and conceptualization (JL, XL, CH, PL, LH), methods development and data interpretation (JL, XL), writing – original draft (JL, XL, LH), and manuscript preparation (JL, XL, CH, PL, LH).

## Figures and Tables

**Fig S1:**
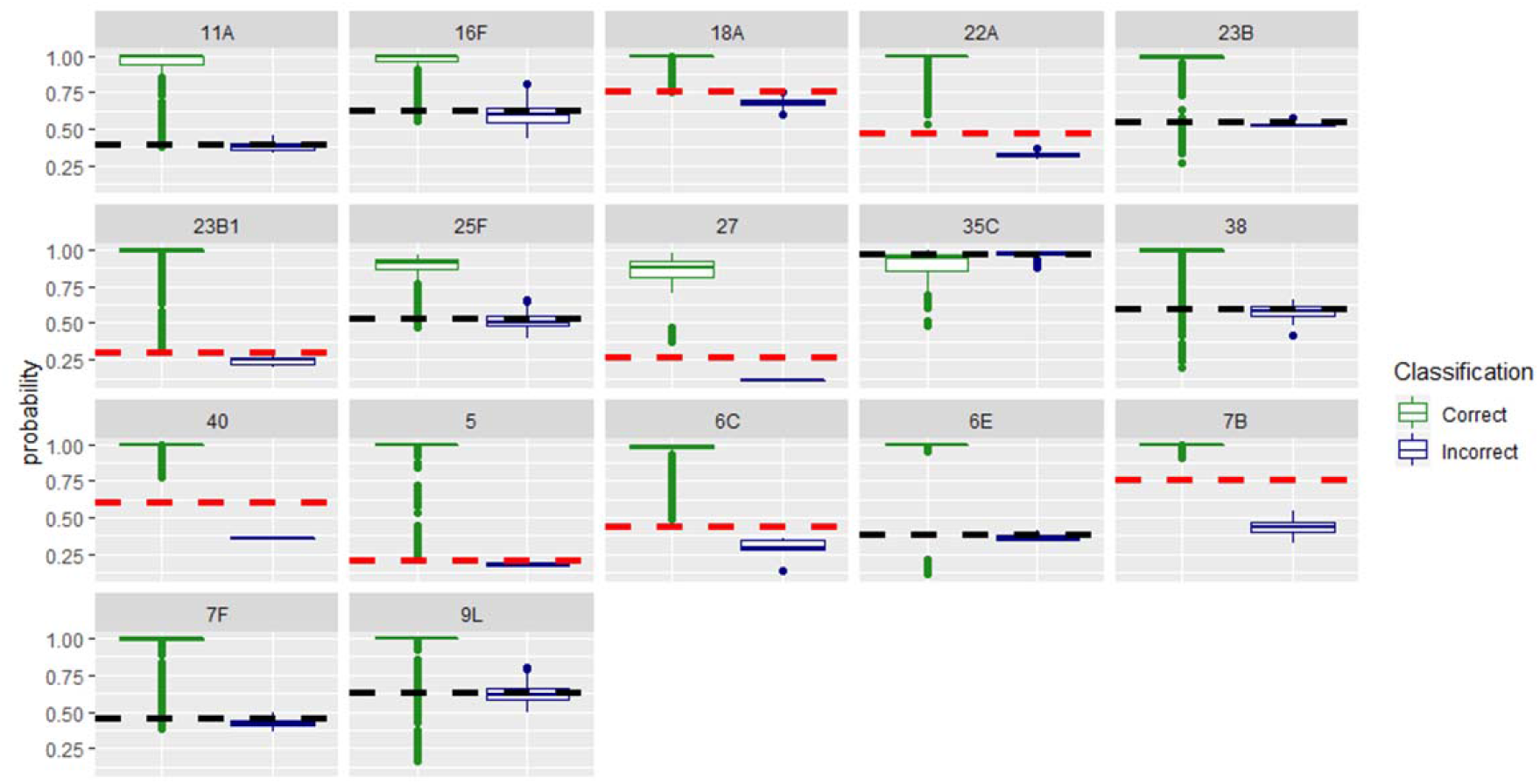
Probability thresholds for 17 serotypes. Model-computed probability distributions for correct and incorrect predictions from cross-validation are shown for 17 serotypes that returned incorrect predictions during model development. Red lines indicate the calculated probability thresholds when distributions do not overlap, and black lines when tails of the distributions do overlap. Predictions made at a probability below these set values are flagged as low-confidence. Calculations for modeling the probability distributions and threshold determination are described in the Materials and Methods.

**Fig S2:**
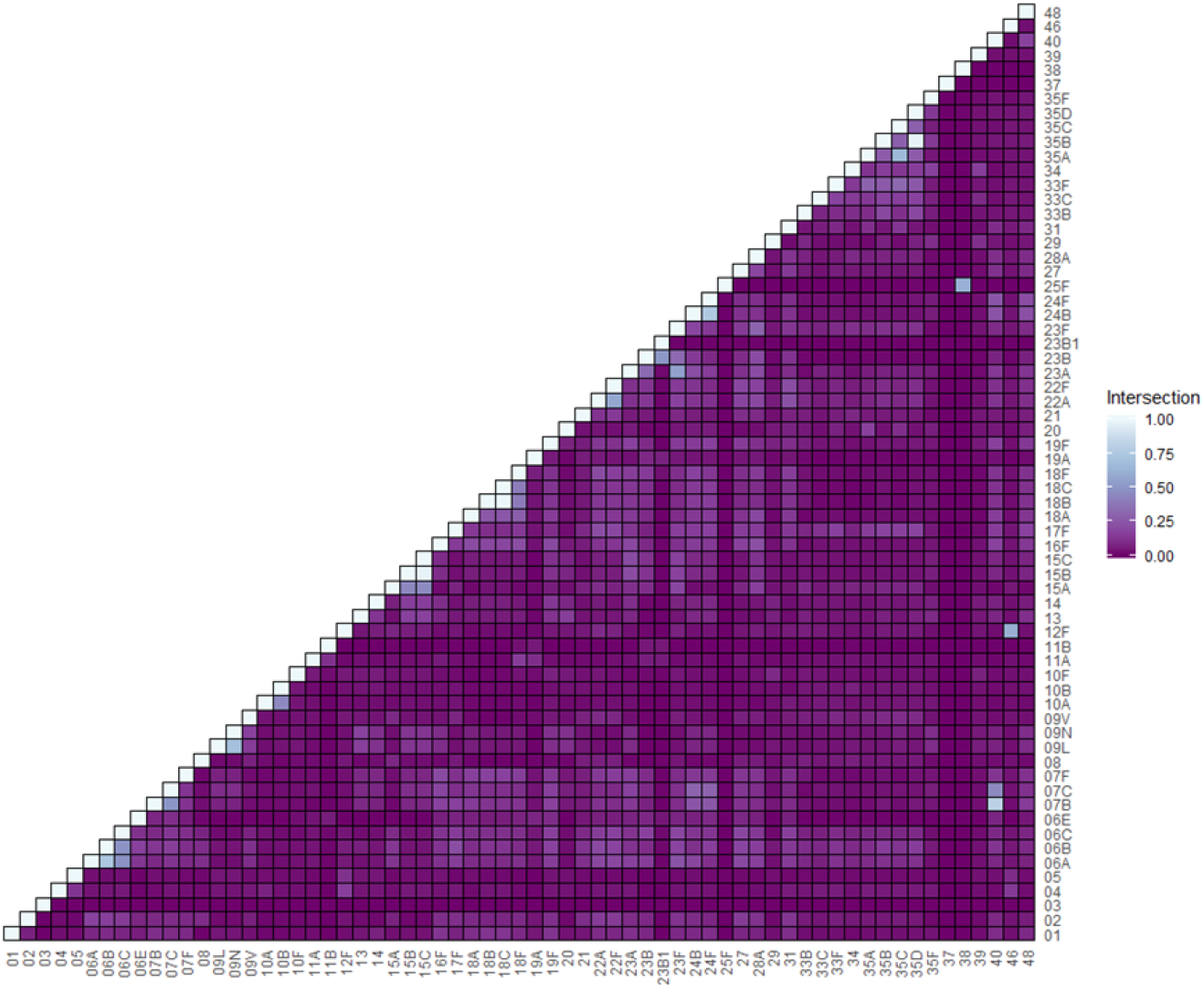
MinHash overlap across serotypes. Fraction of k-mers shared between each pair of 65 serotype MinHash sketches. Proportions correspond to overlap ranging from 0 to 1,000 k-mers.

**Fig S3:**
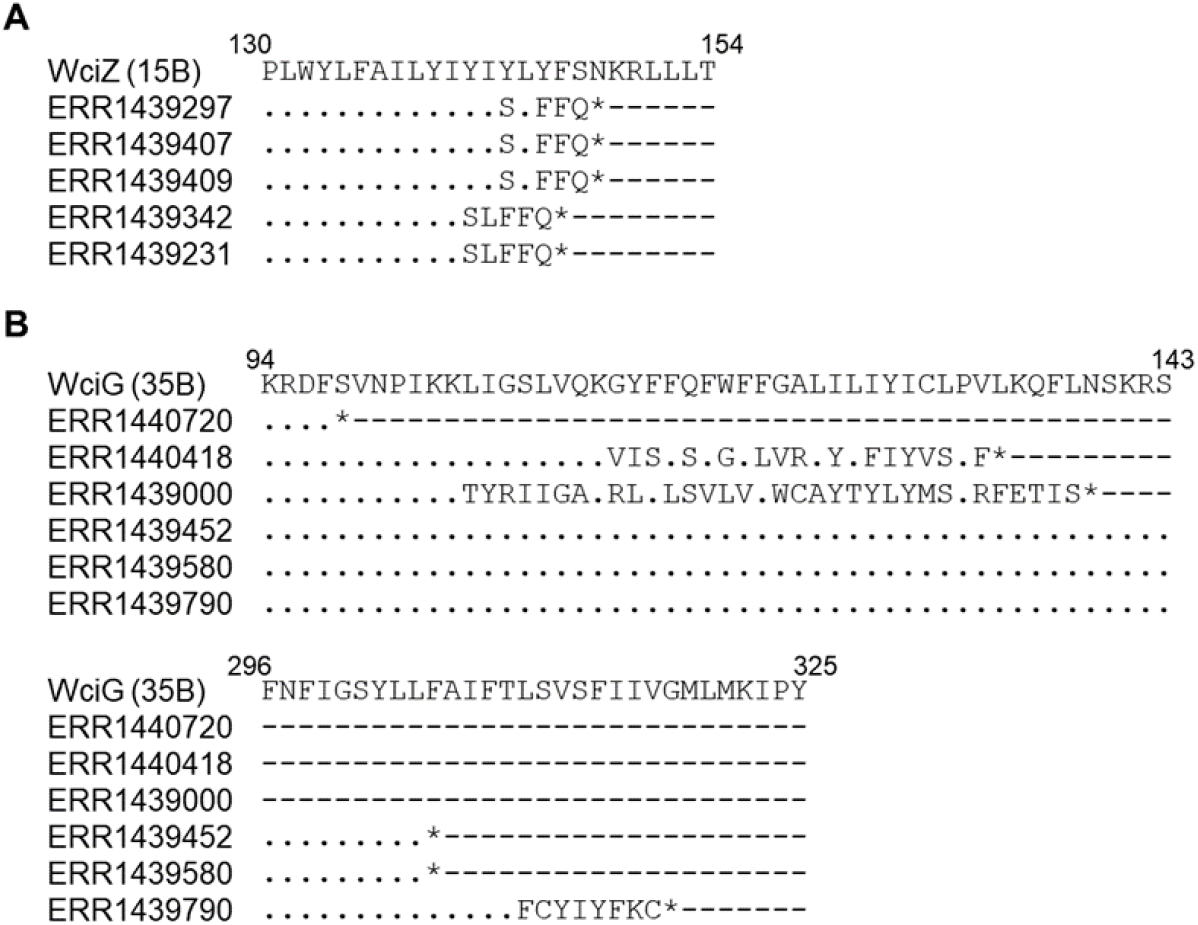
Amino acid sequences of O-acetyltransferases in 15C and 35D isolates predicted by PfaSTer. A) Alignment of a serotype 15B WciZ sequence to five isolates predicted as serotype 15C by PfaSTer. Two variants of WciZ were observed across five isolates, both containing a premature stop prior to residue 150. B) Alignment of a serotype 35B WciG sequence to six isolates predicted to be serotype 35D by PfaSTer. Three sequences terminate prematurely prior to residue 140, and three terminate further downstream prior to residue 320.

**Fig S4:**
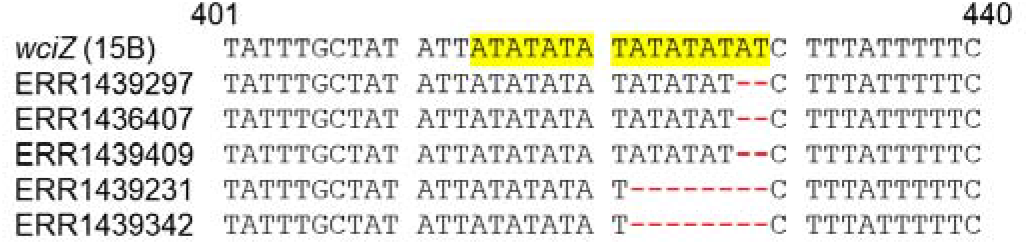
Frameshift mutations at a tandem repeat region of *wciZ* in serotype 15C. Isolates predicted to be serotype 15C by PfaSTer carry multiple nucleotide deletions (red) in an AT-rich tandem repeat (yellow) relative to a 15B reference sequence. Previous work has shown that the resulting frameshift inactivates the WciZ protein and causes formation of the 15C capsular polysaccharide.

**Table S1: Isolate names (pathogen.watch) and serotypes for samples used in PfaSTer training.**

**Table S2: ENA accessions and serotype caller results for isolates used in external validation.**

**Table S3: Fraction of incorrect prediction for each serotype class during cross validation.**

## References

1. Blasi, F., et al., Understanding the burden of pneumococcal disease in adults. Clin Microbiol Infect, 2012. 18 Suppl 5: p. 7–14.

2. Collaborators, G.B.D.L.R.I., Estimates of the global, regional, and national morbidity, mortality, and aetiologies of lower respiratory infections in 195 countries, 1990-2016: a systematic analysis for the Global Burden of Disease Study 2016. Lancet Infect Dis, 2018. 18(11): p. 1191–1210.

3. Drijkoningen, J.J. and G.G. Rohde, Pneumococcal infection in adults: burden of disease. Clin Microbiol Infect, 2014. 20 Suppl 5: p. 45–51.

4. Harboe, Z.B., et al., Impact of 13-valent pneumococcal conjugate vaccination in invasive pneumococcal disease incidence and mortality. Clin Infect Dis, 2014. 59(8): p. 1066–73.

5. McLaughlin, J.M., et al., Effectiveness of 13-Valent Pneumococcal Conjugate Vaccine Against Hospitalization for Community-Acquired Pneumonia in Older US Adults: A Test-Negative Design. Clin Infect Dis, 2018. 67(10): p. 1498–1506.

6. Mavroidi, A., et al., Genetic relatedness of the Streptococcus pneumoniae capsular biosynthetic loci. J Bacteriol, 2007. 189(21): p. 7841–55.

7. Hausdorff, W.P. and W.P. Hanage, Interim results of an ecological experiment - Conjugate vaccination against the pneumococcus and serotype replacement. Hum Vaccin Immunother, 2016. 12(2): p. 358–74.

8. Essink, B., et al., Pivotal Phase 3 Randomized Clinical Trial of the Safety, Tolerability, and Immunogenicity of 20-Valent Pneumococcal Conjugate Vaccine in Adults Aged >/=18 Years. Clin Infect Dis, 2022. 75(3): p. 390–398.

9. Jauneikaite, E., et al., Current methods for capsular typing of Streptococcus pneumoniae. J Microbiol Methods, 2015. 113: p. 41–9.

10. Epping, L., et al., SeroBA: rapid high-throughput serotyping of Streptococcus pneumoniae from whole genome sequence data. Microb Genom, 2018. 4(7).

11. Porter, B.D., B.D. Ortika, and C. Satzke, Capsular Serotyping of Streptococcus pneumoniae by latex agglutination. J Vis Exp, 2014(91): p. 51747.

12. Kapatai, G., et al., Whole genome sequencing of Streptococcus pneumoniae: development, evaluation and verification of targets for serogroup and serotype prediction using an automated pipeline. PeerJ, 2016. 4: p. e2477.

13. Knight, J.R., et al., Determining the serotype composition of mixed samples of pneumococcus using whole-genome sequencing. Microb Genom, 2021. 7(1).

14. Jolley, K.A., J.E. Bray, and M.C.J. Maiden, Open-access bacterial population genomics: BIGSdb software, the PubMLST.org website and their applications. Wellcome Open Res, 2018. 3: p. 124.

15. Ondov, B.D., et al., Mash: fast genome and metagenome distance estimation using MinHash. Genome Biol, 2016. 17(1): p. 132.

16. Ondov, B.D., et al., Mash Screen: high-throughput sequence containment estimation for genome discovery. Genome Biol, 2019. 20(1): p. 232.

17. McGinnis, S. and T.L. Madden, BLAST: at the core of a powerful and diverse set of sequence analysis tools. Nucleic Acids Res, 2004. 32(Web Server issue): p. W20–5.

18. Bankevich, A., et al., SPAdes: a new genome assembly algorithm and its applications to single-cell sequencing. J Comput Biol, 2012. 19(5): p. 455–77.

19. Ganaie, F., et al., Structural, Genetic, and Serological Elucidation of Streptococcus pneumoniae Serogroup 24 Serotypes: Discovery of a New Serotype, 24C, with a Variable Capsule Structure. J Clin Microbiol, 2021. 59(7): p. e0054021.

20. Spencer, B.L., et al., The Pneumococcal Serotype 15C Capsule Is Partially O-Acetylated and Allows for Limited Evasion of 23-Valent Pneumococcal Polysaccharide Vaccine-Elicited Anti-Serotype 15B Antibodies. Clin Vaccine Immunol, 2017. 24(8).

21. McEllistrem, M.C., Genetic diversity of the pneumococcal capsule: implications for molecular-based serotyping. Future Microbiol, 2009. 4(7): p. 857–65.

22. Lo, S.W., et al., Global Distribution of Invasive Serotype 35D Streptococcus pneumoniae Isolates following Introduction of 13-Valent Pneumococcal Conjugate Vaccine. J Clin Microbiol, 2018. 56(7).

23. Malley, J.D., et al., Probability machines: consistent probability estimation using nonparametric learning machines. Methods Inf Med, 2012. 51(1): p. 74–81.

24. van Selm, S., et al., Genetic basis for the structural difference between Streptococcus pneumoniae serotype 15B and 15C capsular polysaccharides. Infect Immun, 2003. 71(11): p. 6192–8.

25. Hao, L., et al., Streptococcus pneumoniae serotype 15B polysaccharide conjugate elicits a cross-functional immune response against serotype 15C but not 15A. Vaccine, 2022. 40(33): p. 4872–4880.

26. Ndlangisa, K., et al., Invasive Disease Caused Simultaneously by Dual Serotypes of Streptococcus pneumoniae. J Clin Microbiol, 2018. 56(1).

27. Zhou, M., et al., Serotype Distribution, Antimicrobial Susceptibility, Multilocus Sequencing Type and Virulence of Invasive Streptococcus pneumoniae in China: A Six-Year Multicenter Study. Front Microbiol, 2021. 12: p. 798750.

28. Ceyhan, M., et al., Serotype distribution of Streptococcus pneumonia in children with invasive disease in Turkey: 2015-2018. Hum Vaccin Immunother, 2020. 16(11): p. 2773–2778.

29. Habibi Ghahfarokhi, S., et al., Serotype Distribution and Antibiotic Susceptibility of Streptococcus pneumoniae Isolates in Tehran, Iran: A Surveillance Study. Infect Drug Resist, 2020. 13: p. 333–340.

30. Isturiz, R., et al., Expanded Analysis of 20 Pneumococcal Serotypes Associated With Radiographically Confirmed Community-acquired Pneumonia in Hospitalized US Adults. Clin Infect Dis, 2021. 73(7): p. 1216–1222.

31. Lister, A.J.J., et al., Serotype distribution of invasive, non-invasive and carried Streptococcus pneumoniae in Malaysia: a meta-analysis. Pneumonia (Nathan), 2021. 13(1): p. 9.

32. Wiese, A.D., M.R. Griffin, and C.G. Grijalva, Impact of pneumococcal conjugate vaccines on hospitalizations for pneumonia in the United States. Expert Rev Vaccines, 2019. 18(4): p. 327–341.

