## Supplemental Note S1 for "PfaSTer: A ML-powered serotype caller for *Streptococcus pneumoniae* genomes"

Serotype 23B1 was not included in agglutination assays – instead labeled 23B, but was recognized by both PneumoCaT and SeroBA. In these instances, PfaSTer predictions were considered in concordance when also returning 23B1. Serotype 35D was not supported by SeroBA v0.0.1, for which results were initially published. Samples labeled 35B were reanalyzed with SeroBA v1.0.1 and cases where 35D was reported are included in Table S2. Additionally, neither the published results for SeroBA nor PfaSTer designate 6A/6E and 6B/6E, so the subtypes 6A/6B and 6E were considered in concordance when predicted by these tools. In cases where latex agglutination or another in-silico method returned multiple serotypes (or an entire serogroup), all the serotypes listed were taken into account during comparison to PfaSTer.
